# Protective role of the deSUMOylating enzyme SENP3 in myocardial ischaemia-reperfusion injury

**DOI:** 10.1101/557249

**Authors:** Nadiia Rawlings, Laura Lee, Yasuko Nakamura, Kevin A. Wilkinson, Jeremy M Henley

**Affiliations:** School of Biochemistry, Centre for Synaptic Plasticity, Biomedical Sciences Building, University of Bristol, University Walk, Bristol, BS8 1TD, U.K.

## Abstract

Interruption of blood supply to the heart is a leading cause of death and disability. However, the molecular events that occur during heart ischaemia, and how these changes prime consequent cell death upon reperfusion, are poorly understood. Protein SUMOylation is a post-translational modification that has been strongly implicated in the protection of cells against a variety of stressors, including ischaemia-reperfusion. In particular, the SUMO2/3-specific protease SENP3 has emerged as an important determinant of cell survival after ischaemic infarct. Here, we used the Langendorff perfusion model to examine changes in the levels and localisation of SUMOylated target proteins and SENP3 in whole heart. We observed a 50% loss of SENP3 from the cytosolic fraction of hearts after preconditioning, a 90% loss after ischaemia and an 80% loss after ischaemia-reperfusion. To examine these effects further, we performed ischaemia and ischaemia-reperfusion experiments in the cardiomyocyte H9C2 cell line. Similar to whole hearts, ischaemia induced a decrease in cytosolic SENP3. Furthermore, shRNA-mediated knockdown of SENP3 led to an increase in the rate of cell death upon reperfusion. Together, our results indicate that cardiac ischaemia dramatically alter levels of SENP3 and suggest that this may a mechanism to promote cell survival after ischaemia-reperfusion in heart.

## Introduction

Standard clinical treatment after a heart attack is to restore blood supply as soon as possible in order to limit infarct size and reduce mortality. Paradoxically, however, this can cause oxidative damage, referred to as reperfusion injury, which leads to cardiomyocyte cell death and contributes to reduced cardiac output (1). An important goal in the field is to identify the cellular changes that occur during ischaemia and subsequent reperfusion, with a view to determining how intervening in these pathways may promote cell viability after ischaemic insult.

Particularly relevant is the phenomenon of ischaemic preconditioning, whereby tissues exposed to repeated short bursts of ischaemia protect the tissue against the effects of prolonged ischaemia. How preconditioning leads to cellular protection is poorly understood so defining the pathways involved and designing strategies to potentiate them to promote cellular survival after ischaemia represent important goals. Important clues come from animals that hibernate because they have very low rates of blood flow, so tissues endure prolonged hypoxia but emerge from hibernation torpor undamaged. It has emerged that one protective mechanism is massively increased levels of protein post-translational modification of lysine residues by small ubiquitin-like modifier (SUMO) during torpor (2, 3).

There are three ∼11 kDa SUMO paralogues (SUMO-1-3) in mammals that are conjugated to substrates, generally within a consensus motif (4, 5). While the functional consequences of SUMOylation vary depending on the substrate, the underlying principle is that it alters inter- and/or intramolecular interactions of substrate proteins to change their localisation, stability, and/or activity (6).

SUMOylation is dynamic and SUMO can be removed from proteins by SUMO proteases (SPs). Nine mammalian SPs have been identified but their specific targets and physiological roles are poorly characterized and how SP activity is regulated to control substrate deSUMOylation is largely unknown (reviewed in (7-9)). The largest family of SPs are the sentrin-specific proteases (SENPs). Generally, SENP1 and SENP2 mature pre-SUMOs and deconjugate SUMO1 and SUMO2/3, SENP3 and SENP5 preferentially deconjugate SUMO2/3, and SENP6 and SENP7 can edit polySUMO2/3 chains (10, 11).

Significant changes in SUMO pathway components occur during ischaemia in the human heart (12). Moreover, over-expression of SUMO enhances cardiac function in mice with heart failure and increases contractility in isolated cardiomyocytes (13). Interestingly, recent data also suggests a role for SENP3 in determining cardiomyocyte survival after ischaemia-reperfusion, however whether SENP3 promotes cell survival or cell death remains controversial. It has been reported that SENP3 levels are upregulated in heart tissue in response to ischaemia-reperfusion, and that knockdown of SENP3 *in vivo* reduces infarct size and improves cardiac function (14). Similarly, in the cardiomyocyte H9C2 cell line, there is an increase in SENP3 levels in response to ischaemia-reperfusion but, in contrast, knockdown of SENP3 promoted apoptosis after ischaemia-reperfusion (15).

To directly address this controversy and characterise the molecular events that occur during ischaemia and reperfusion, we investigated how SUMOylation and levels of SENP3 are altered by ischaemia and ischaemia-reperfusion in whole heart. To investigate the molecular mechanisms that underlie cell protection afforded by ischaemic preconditioning, we also examined hearts which were subjected to 3 short bursts of ischaemia-reperfusion prior to the extended ischaemic insult. Cytosolic levels of SENP3 were dramatically reduced by ischaemia and ischaemia-reperfusion. Furthermore, knockdown of SENP3 in the cardiomyocyte H9C2 cell line demonstrated that loss of SENP3 increases the rate of cell death in response to ischaemia-reperfusion. Together, these data suggest an important role for SENP3-mediated deSUMOylation in promoting cardiomyocyte survival after ischaemic insult.

## Materials and methods

### Heart perfusion

All procedures were performed in accordance with Schedule 1 Guidance on the Operation of the Animals (Scientific Procedures) Act 1986 and are described previously (16). In brief, male Wistar rats (250-275g) were anaesthetised with isofluorane and killed by cervical dislocation. Hearts were rapidly cannulated via the aorta *in vivo*, then quickly removed and placed into 37°C buffered Krebs-Henseleit solution containing in (mM): NaCl 118, NaHCO_3_ 25, KCl 4.8, KH_2_PO_4_ 1.2, MgSO_4_ 1.2, glucose 11 and CaCl_2_ 1.2, gassed with 5% CO_2_ at 37°C (pH 7.4). Hearts were perfused at 12 ml min^-1^ in the Langendorff mode with in-line filter using Krebs-Henseleit buffer. No electrical stimulation was applied. Left ventricular developed pressure (LVDP) was monitored with a water-filled balloon inserted into the left ventricle. Initial end diastolic pressure (EDP) was set to give an initial value of 4–5 mm Hg. All hearts were allowed an equilibration period of at least 20 min before the time for each group was started. Control group (CTL) was perfused for 45 min. Global normothermic ischaemia was induced by switching off the pump and submersing the heart in buffer kept at 37°C. In all groups the ischaemic period was 30 min. The preconditioned-ischaemic group was subjected to three cycles of 2 min of global ischaemia interspersed with 3 min reperfusion prior to 30 min global normothermic ischaemia. In the ischaemia-reperfusion group, after 30 min of global ischaemia buffer flow was restored for the next 2 hours.

### Subcellular fractionation of left ventricular tissue

Isolation of subcellular fractions from rat left ventricular tissue was adapted from a previously published protocol (17). Left ventricular tissue was immersed in 6 ml ice cold isolation buffer (300 mM sucrose, 3 mM EGTA, 10 mM Tris-HCl, pH 7.1) supplemented with 20 mM NEM, 1x complete protease inhibitors (Roche) and 1x phosphatase inhibitors (Roche). Tissue was rapidly chopped with scissors into fine pieces before homogenisation using a Polytron tissue disruptor (Kinematica) at 10,000rpm with 2 bursts of 5 seconds followed by 1 burst of 10 seconds. Tissue homogenate was diluted to 20 ml total volume with isolation buffer supplemented with 1x complete protease inhibitors and further homogenised by hand for 2 minutes using a glass Potter homogeniser and Teflon pestle. A small volume of homogenate was stored at −80°C as whole homogenate. The rest of the homogenate was centrifuged at 7500g for 7 minutes and the soluble fraction (supernatant) stored at −80°C as a cytosol-containing fraction. The pellet was rinsed twice with 5 ml isolation buffer, resuspended in 20mL isolation buffer and further hand-homogenised for 2 minutes. The homogenate was then centrifuged at 600g for 10 minutes and the pellet resuspended in isolation buffer and stored at −80°C as a nucleus-containing fraction. The supernatant was centrifuged at 7000g for 10 minutes to yield a crude mitochondrial pellet, which was resuspended in isolation buffer and stored at −80°C. Proteins from soluble cytosol-containing fraction were precipitated with acetone for at least 1h at −20°C, spun down and resuspended in isolation buffer. All fractions were assayed for protein concentration using a standard BCA assay protocol prior to Western blotting.

### Senp3 knockdown

An shRNA sequence targeting rat SENP3 (target sequence TATGGACAGAACTGGCTCAATGACCAGGT), or a non-targeting (‘scrambled’) control (target sequence GCACTACCAGAGCTAACTCAGATAGTACT) were cloned into a modified pXLG viral vector under the control of a U6 promoter. Viral particles were produced in HEK293T cells using the helper vectors pMD2.G and p8.91 as described previously (18), and virus-containing supernatant used to transduce H9C2 cells.

### Lactate dehydrogenase (LDH) assay

Medium was collected from experimental wells, placed on ice, spun down at 1000g for 1 min and tested for LDH within 30 min of collection. The LDH cytotoxicity detection kit was purchased from Clontech and used according to the manufacturer’s protocol. Briefly, one volume each of tested sample and working solution were mixed in a 96 well plate, and incubated for 30 minutes at RT, protected from light. The absorbance was then read in a spectrophotometer at 490nm and 620nm. LDH activity in each well was calculated by subtracting the 620nm reading (plastic plate background) from the 490nm reading. Each experiment contained the following controls: background control (medium only) – subtracted from all values as a background reading; high control - measures the maximum LDH activity that can be released from the 100% dead cells in response to medium containing 1% Triton X-100. Values of % dead cells in experimental wells were calculated with respect to the 100% dead cells value from Triton X-100 treated cells. Importantly, for these experiments, media lacking phenol red and sodium pyruvate was used as both components interfere with LDH readings.

### H9C2 cell culture and ischaemia-reperfusion protocol

H9C2 cells were cultured in media (FG 0435, Biochrome) containing 10% FBS Superior (S0615, Biochrome) and 0.05% penicillin/streptomycin for 5 days. For LDH measurements 20×10^3^ cells were plated per well in a 6-well plate; for OGD experiments – 120×10^3^ cells per 100 mm dish. 2 days before experiments 10% FBS-containing medium was replaced with medium containing 1% horse serum (New Zealand origin, Invitrogen) and 0.05% penicillin/streptomycin.

For ischaemia only or ischaemia-reperfusion experiments all cells were washed with PBS and changed from phenol red-containing culture medium to transparent media (Gibco, A1443001) containing 1% horse serum and 4.5g/L glucose for 1h. All time-matched controls had their glucose-containing media replaced with the same glucose-containing media the same amount of times as their respective ischaemia or ischaemia-reperfusion conditions. Ischaemia was achieved by placing cells in the MACS-VA500 anaerobic workstation (Don Whitley Scientific limited) supplemented with 95% N_2_ and 5% CO_2_ at 37°C. Once in the anaerobic workstation, cells were washed with deoxygenated PBS and then placed in deoxygenated medium containing 1% horse serum and lacking glucose for the specified time. In order to achieve quick reperfusion, at the end of ischaemia, cells were quickly changed back into oxygen and glucose-containing media inside the chamber and plates were placed back in standard cell culture incubators (5% CO_2_, 37°C).

### H9C2 subcellular fractionation

Before collection all cells were washed 4 times with ice-cold PBS to remove horse serum-containing media. Subcellular fractionation was then performed based on a previously published protocol (19) with some modifications. Briefly, cells from a 100mm dish were collected in 1 ml of prechilled lysis buffer A (100 mM NaCl in 50 mM HEPES containing 25 μg/ml digitonin, protease and phosphatase inhibitor cocktails and 20 mM NEM). Cells were then hand homogenized in a glass homogenizer on ice and spun for 30 min at 16000g at 4°C. Supernatant was collected as a cytosolic sample, and pellets were collected as a membrane/nuclear fraction. The membrane/nuclear-containing pellet was then resuspended and sonicated in lysis buffer B (100 mM NaCl in 50 mM HEPES buffer, containing 5 mg /ml sodium deoxycholate, 1% Triton-X, 0.5% SDS, protease and phosphatase inhibitor cocktails and 20 mM NEM). Since the cytosolic fraction was too dilute, proteins were precipitated in 5 volumes of prechilled acetone for 2h at −20°C, then spun down for 30 min at 3000g at 4°C. The pellet containing the cytosolic sample was then resuspended and sonicated in lysis buffer B. All samples were evaluated for protein concentration by BCA assay (Thermo Fisher).

### Immunoblotting

Samples were resolved by SDS-PAGE (7.5-15% gels), transferred to Immobilon-P membranes (Millipore Inc.) and immunoblotted as indicated. Primary antibodies used were: SUMO-1 and SUMO-2/3 (sheep, a gift from Ron Hay (University of Dundee)), VDAC (Cell Signalling, D73D12), GAPDH (Sigma, G9295), SENP3 (Cell Signaling, D20A10), Lamin (Santa Cruz, sc-6216), and cleaved caspase 3 (Cell Signaling, 5A1E). Total protein on membranes was stained using REVERT total protein stain (LI-COR). Immune complexes were detected using HRP-conjugated secondary antibodies (Sigma) followed by enhanced chemiluminescence (Thermo Scientific Pierce) using a LI-COR Odyssey machine or x-ray films. Each immunoblot presented is representative of at least three experiments carried out using different cell populations.

### Statistics

All statistical analyses are specified in the Figure legends.

## Results

### Characterisation of haemodynamic properties of perfused hearts

To examine the molecular changes that occur during ischaemia, ischaemia-reperfusion and ischaemic-preconditioning, we used the Langendorff perfused isolated rat heart model. We chose to evaluate 4 experimental groups:

i. **Control**, 20 min of stabilization perfusion and 50 min perfusion with Krebs-Henseleit buffer (KHB).
ii. **Preconditioning and ischaemia (PCI)**, 20 min of stabilization perfusion followed by 3 cycles of 2 min ischaemia and 3 min reperfusion (15 min of preconditioning) and then 30 min of ischaemia.
iii. **Ischaemia**, 20 min of stabilization perfusion, 20 min perfusion and 30 min of global ischaemia.
iv. **Ischaemia + reperfusion (I/R)**, 20 min of stabilization perfusion, 20 min perfusion, 30 min global ischaemia and 2h of reperfusion.

We first examined several haemodynamic parameters to validate the model. These were left ventricular developed pressure (LVDP), end-diastolic pressure (EDP), heart rate, time to ischaemic contracture and maximal contracture. As shown in Table 1, in preconditioned hearts the time to ischaemic contracture was significantly shorter than in ischaemic hearts, indicating preconditioning occurred in our hearts consistent with previous observations (20). Moreover, since time to ischaemic contracture was the same in the ischaemia and I/R groups, we conclude that there was no unintended preconditioning in the I/R group and that any differences between these groups can be attributed to ischaemia-reperfusion injury.

**Table 1.** Haemodynamic parameters for Langendorff perfused hearts.

|  | Control<br>n=6 | Preconditioned-Ischaemia<br>n=5 | Ischaemia<br>n=6 | Ischaemia /<br>reperfusion<br>n=6 |
| --- | --- | --- | --- | --- |
| *20 min point* |  |  |  |  |
| LVPD (mm Hg) | 79.7±14.4 | 81.8±3.6 | 73.3±8.4 | 76.0±5.6 |
| EDP (mm Hg) | 8.4±0.8 | 8.0±1.0 | 11.3±1.2 | 7.6±0.9 |
| HR (bpm) | 321±7 | 308±13 | 310±13 | 338±19 |
| *Ischaemia* |  |  |  |  |
| Time to IC (min) | - | 7.7±1.2** | 13.5±0.8 | 12.8±0.5 |
| Max IC (mm Hg) | - | 68.4±10.0 | 54.6±7.0 | 42.4±3.9 |

All values are given as mean ± SEM. Data are shown for hearts used for further sample collection and Western blot sampling. LVPD – left ventricular developed pressure; EDP – end-diastolic pressure; HR – heart rate; IC – ischaemic contracture. Note that because we collected our heart samples for biochemical analyses at the end of each described stage, we were unable measure recovery of LVPD for the preconditioned group since there was no reperfusion after ischaemia. \*\* Significant difference between PCI group and Isc and I/R groups (p<0.01). One-way ANOVA with Bonferroni's multiple comparison test was used.

### Subcellular fractionation of whole hearts

To profile changes in SUMOylation and SENP3, we prepared whole tissue homogenates from hearts that had undergone the Langendorff protocol as well as separate cytosolic, nuclear and mitochondrial fractions. Following trial experiments to optimise the fraction separation and retain mitochondria-associated proteins we used a relatively gentle mitochondrial enrichment fractionation protocol (see methods).

### Protein SUMOylation in perfused heart

Having validated the experimental system, we next measured levels of SUMO1- and SUMO2/3-ylation in total homogenates and in each subcellular fraction from the perfused hearts. No significant changes were detected in total SUMO-immunoreactivity in whole homogenates (Fig. 1A). Similarly, we did not detect any overall changes in protein SUMOylation in the mitochondrial fraction (Fig. 1B). It should be noted, however, that these global protein assays do not exclude the possibility of changes in the SUMOylation status of individual or subsets of mitochondrial proteins.

**Fig. 1.**
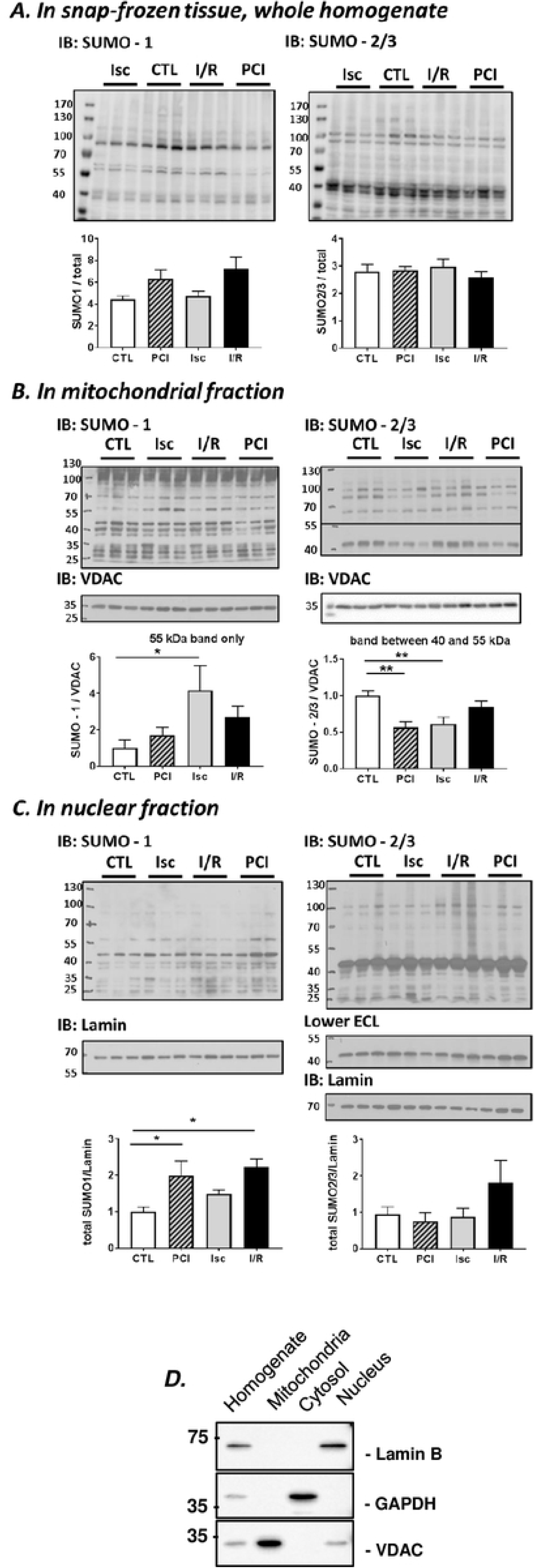
Total SUMOylation of proteins in rat heart during preconditioning-ischaemia (PCI), ischaemia (Isc) only and ischaemia followed by reperfusion (I/R). (**A**) Snap-frozen whole homogenate samples, (**B**) mitochondria-enriched fraction and (**C**) nuclear fractions of perfused left ventricular tissue were Western blotted with SUMO-1 (left) or SUMO-2/3 (right) antibodies. Representative blots are of 3 samples per condition. Total protein stain with Ponceau, or staining with VDAC or Lamin antibodies, were used as a loading control for whole homogenate samples or mitochondrial or nuclear factions, respectively. Quantitative analysis using ordinary one-way ANOVA with Sidak’s correction for multiple comparisons with a pooled variance. N=6 for CTL, PCI and I/R; N=5 for Isc group. Data presented as mean ± SEM. *p<0.05; ** p<0.01. (**D**) Validation of the fractionation protocol used. Equal amounts of whole homogenate or mitochondrial, cytosolic or nuclear fraction from left ventricular rat heart tissue were Western blotted as indicated. Lamin was used as a nuclear marker, GAPDH as a cytosolic marker and VDAC as a mitochondrial marker.

Indeed, an approximately 55kDa SUMO-1-ylated species was significantly upregulated in the mitochondrial fraction under ischaemia, compared to pre-ischaemia, but was not upregulated during ischaemia in tissue that had been preconditioned or after reperfusion. Similarly, while there were no global alterations in SUMO-2/3 conjugation, the SUMOylation of an individual mitochondrial substrate at ∼45kDa was reduced by preconditioning and ischaemia (Fig. 1B).

Similarly, we did not detect changes in total nuclear protein SUMO-2/3-ylation in any of the conditions. In contrast, however, we observed robust changes in SUMO-1-ylation of nuclear proteins, with significantly increased levels in preconditioned and I/R conditions (Fig. 1C). Thus, while no striking differences in the overall levels of SUMOylated proteins in whole homogenate or mitochondria were detected, changes were observed for individual SUMO-1 and SUMO-2/3 substrate proteins, and for total SUMO-1-ylation in the nucleus. The purity of the subcellular fractions were validated using specific mitochondrial, cytosolic and nuclear marker proteins, confirming the high levels of separation of each fraction (Fig. 1D).

### Changes in SENP3 during ischaemia and reperfusion

The extent and duration of substrate SUMOylation is determined by the balance between conjugation and SUMO-protease-mediated deconjugation. SENP3 is one of the best characterised SUMO-proteases, and preferentially removes SUMO-2/3 from target proteins (9). Importantly, SENP3 has recently emerged as a key determinant of cell survival after ischaemic stress (6, 21, 22). We therefore tested the effects of preconditioning, ischaemia and reperfusion on SENP3 levels in total homogenate, cytosol and nuclear fractions of Langendorff perfused whole heart (Fig. 2). There were no significant changes in the total levels of SENP3 between the control, preconditioning, ischaemia and I/R conditions in whole homogenates. In the cytosolic compartment, however, SENP3 levels were significantly decreased in the preconditioned, ischaemia, and I/R groups. Moreover, this loss of SENP3 levels in cytosol during ischaemia was significantly attenuated by prior preconditioning. Correlating with these data, we observed an increase in SENP3 in the nuclear fraction in the ischaemia group, the condition in which there was the biggest decrease in cytosolic SENP3. Taken together, these data suggest that ischaemia leads to a translocation of SENP3 from the cytosol to the nucleus, and that the magnitude of this translocation can be reduced by preconditioning prior to the insult.

**Fig. 2.**
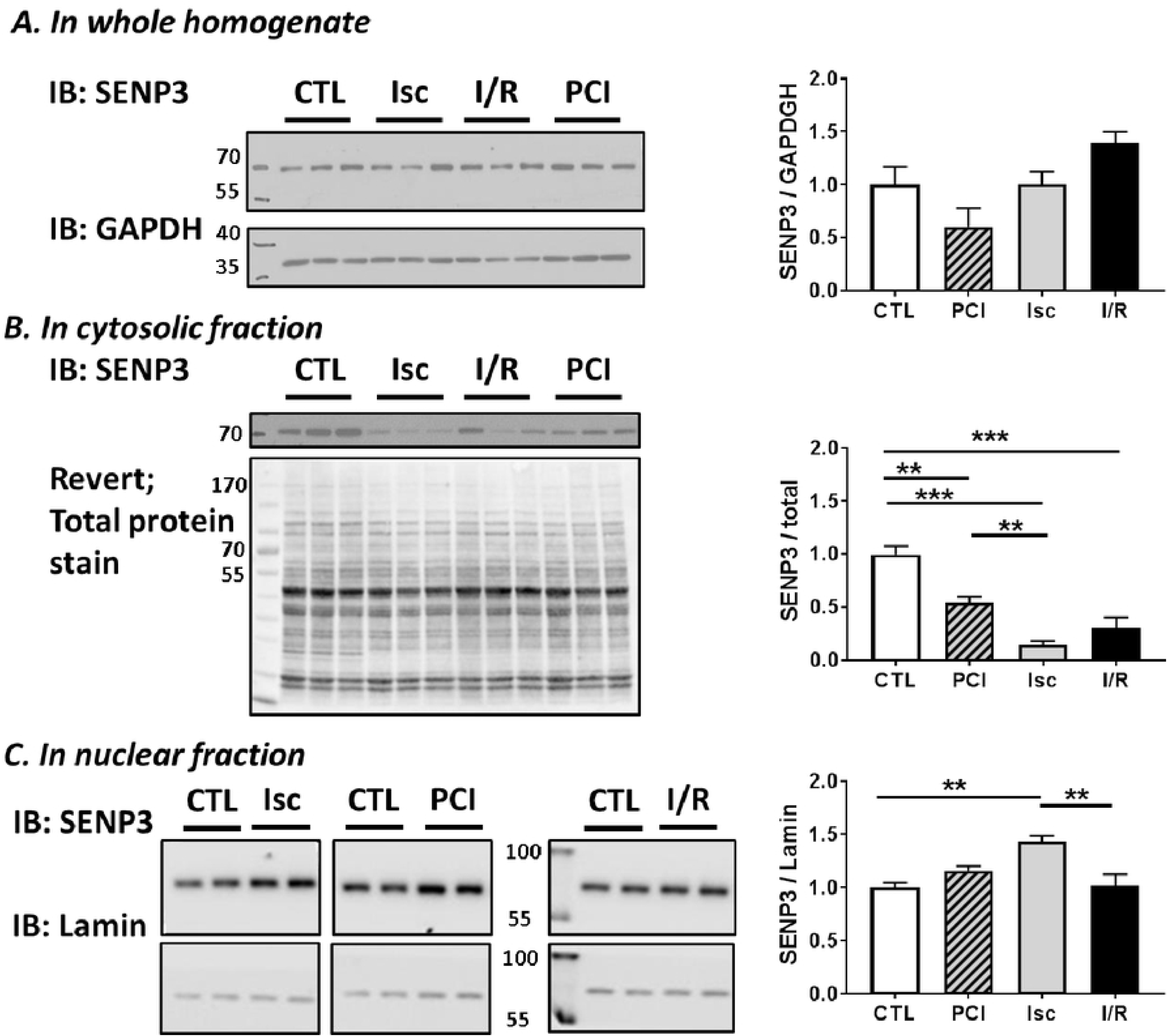
SENP3 levels in different subcellular fractions of rat heart. (**A**) Whole homogenate samples, (**B**) cytosolic and (**C**) nuclear fractions of perfused left ventricular tissue were Western blotted with an antibody against SENP3. Representative blots are of 2-3 samples per condition. REVERT total protein stain or GAPDH or Lamin antibodies were used as loading controls. Quantitative analysis was performed with an ordinary one-way ANOVA with Sidak’s correction for multiple comparisons with a pooled variance. N=6 for CTL, PCI and I/R; N=5 for Isc group. Data presented as mean ± SEM. *p<0.05; ** p<0.01, *** p<0.001.

### Modelling of ischaemia-reperfusion injury in H9C2 cells

Our data from the whole heart ischaemic model show that cytosolic SENP3 levels are reduced during ischaemia (Fig. 2B). We therefore wondered how SENP3 levels affect cell survival in cardiomyocytes in response to ischaemia and ischaemia-reperfusion. To do this, we used the well characterised rat ventricle origin cardiomyoblast H9C2 cell line (23), since this model system allowed us to genetically manipulate SENP3 levels prior to ischaemic insult.

First, we confirmed that H9C2 cells respond to ischaemic insult in a similar way to whole heart, with a significant decrease in cytosolic SENP3 within the first 30 min of ischaemia (a drop of 42±12.3% compared to the control group, p=0.02, t-test, n=3), confirming that this cell line represents a useful model system for examining the role of SENP3 in the cellular response to ischaemic stress.

We next profiled the time course of the response of H9C2 cells to ischaemia to identify appropriate timepoints to examine the role of SENP3 in cell survival. We subjected cells to varying periods of ischaemia and assayed the initiation of apoptosis using cleavage of the executioner caspase 3 (24) as a marker (Fig. 3A). Caspase 3 activation showed a bell-shaped response plotted against time of ischaemia, with the peak at 2h. Additionally, we assayed cell death by measuring LDH release into the culture medium. Interestingly, H9C2 cells appear very resilient to these challenges, requiring 24h of ischaemia alone (Fig. 3B) or minimum 12h of ischaemia followed by 24h of reperfusion (Fig. 3C) to induce significant levels of cell death. Nonetheless, these experiments allowed us to identify a time window in which to examine the role of SENP3 in H9C2 cell survival after ischaemia-reperfusion.

**Fig. 3.**
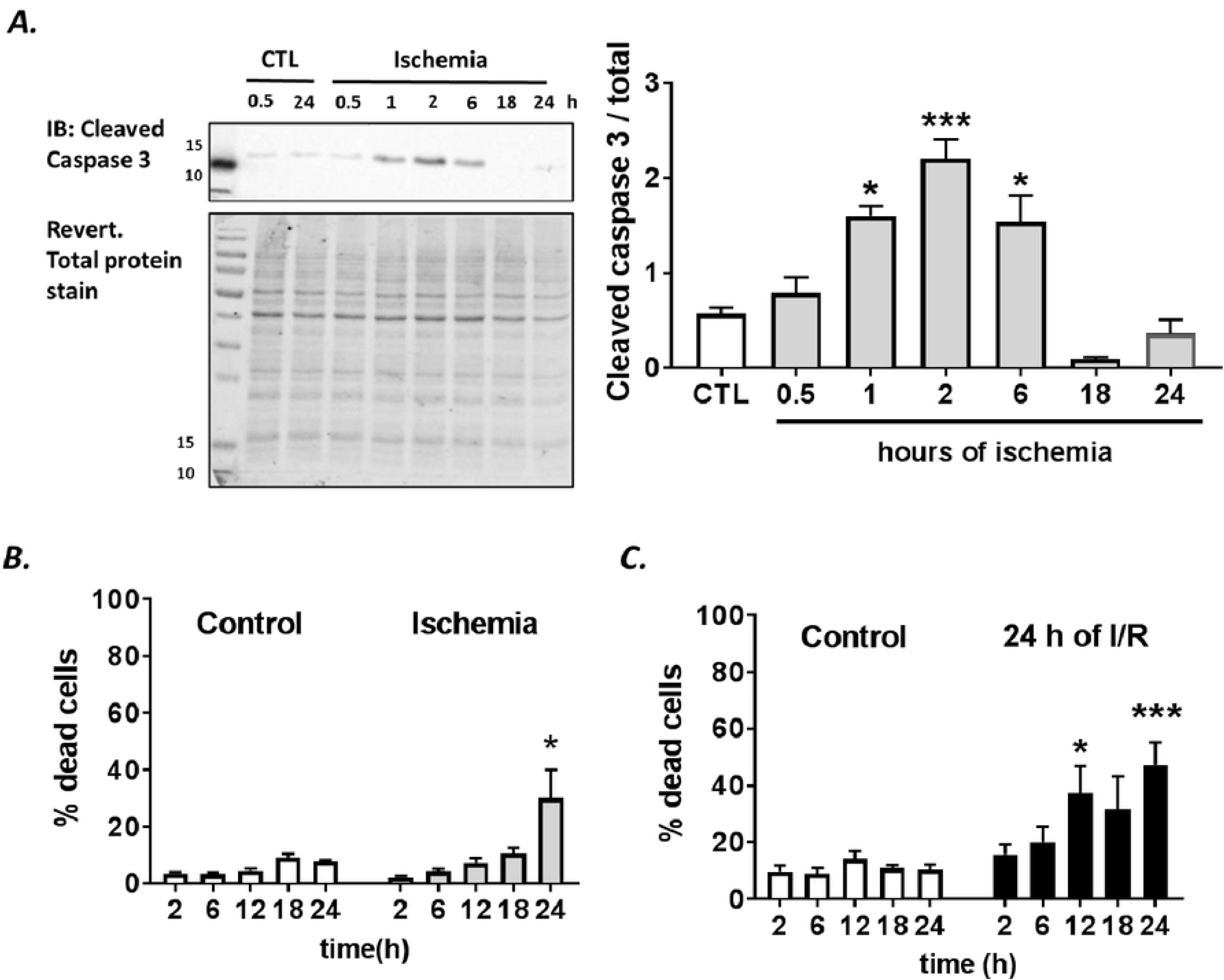
Effect of different lengths of ischaemia on cell death parameters in H9C2 cells. (**A**) Activation of cleaved caspase 3, as monitored by the ∼15kDa cleavage fragment. The signal peaks following 2h of ischaemia. Quantitative analysis using ordinary one-way ANOVA with Sidak’s correction for multiple comparisons with a pooled variance. Data presented as mean ± SEM. *p<0.05; **** p<0.001. (**B**) Profile LDH release assay in ischaemia experiments, N=3. (**C**) Profile of LDH release assay in experiments where 2 −24h ischaemia was followed by 24h of reperfusion, N=3 experiments. For (**B**) and (**C**), Students t-test comparing control to ischaemia at each time point. Data presented as mean ± SEM. *p<0.05; *** p<0.001.

### Effects of SENP3 knockdown on H9C2 cell response to ischaemic stress

Having established conditions to monitor ischaemia-induced cell death in H2C9 cells we next investigated the effects partial SENP3 knockdown. We infected H9C2 cells with lentivirus expressing either a control scrambled non-targeting shRNA or an shRNA targeting SENP3, which reduced levels of SENP3 by ∼50%. 14 DIV after infection cells were subjected to 30 min, 2h or 18h of ischaemia (Fig. 4).

**Fig. 4.**
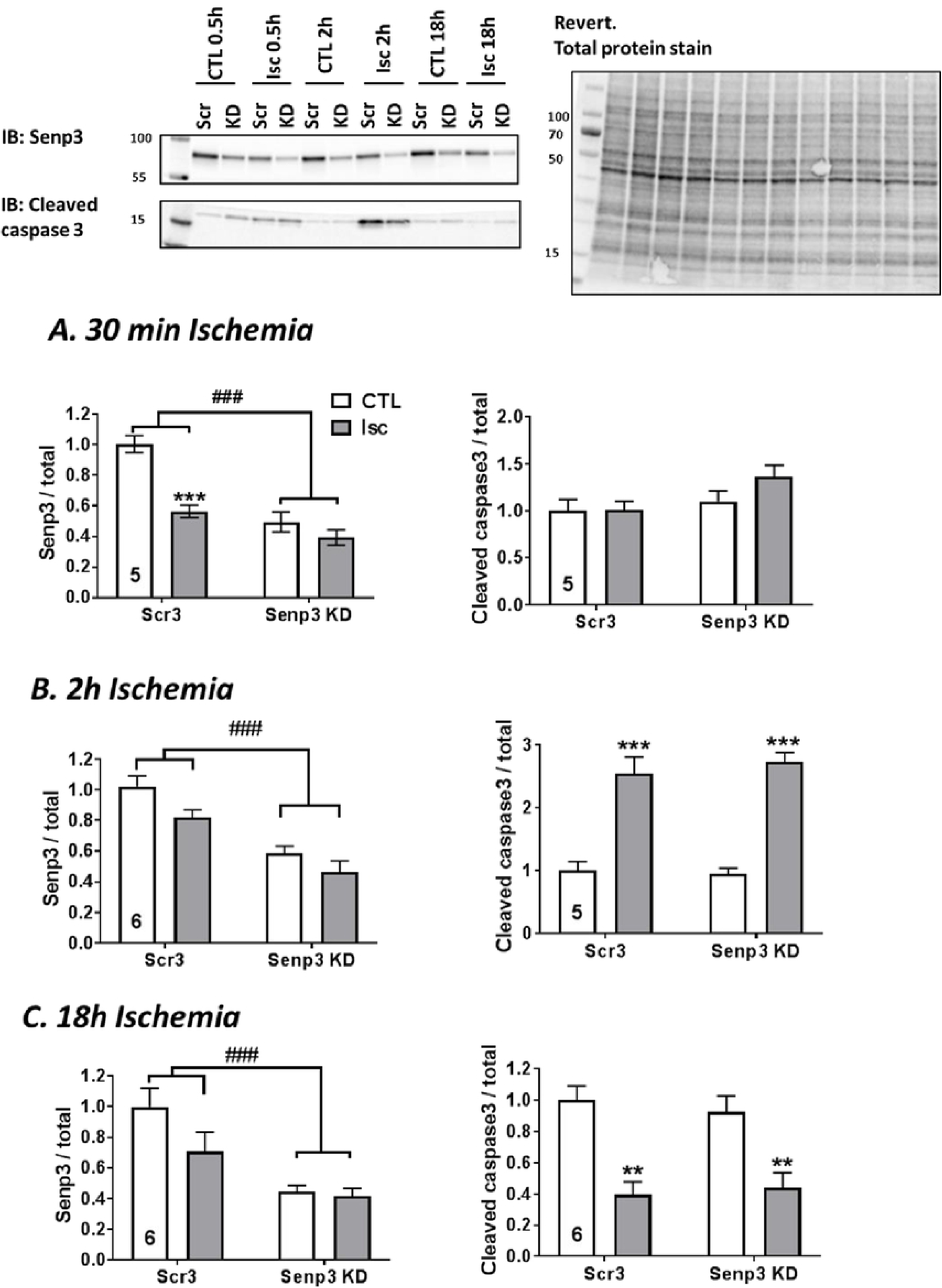
Effects of 30 min, 2h and 18h ischaemia on SENP3 levels and caspase 3 activation in control and SENP3 knockdown H9C2 cells. Western blots of representative experiments. Scr3 = control, scrambled non-targeting shRNA, Senp3 KD = shRNA targeting SENP3. (**A**) 30 min of ischaemia; (**B**) 2h of ischaemia; (**C**) 18h of ischaemia. Quantitative analysis using ordinary two-way ANOVA test with Sidak’s correction for multiple comparisons for levels of SENP3 and cleaved caspase 3. Number of experiments indicated in each graph. Data presented as mean ± SEM. * show p values for effect of ischaemia; # show p values for effect of SENP3 KD.

In the scrambled shRNA control condition, 30 min ischaemia reduced cytosolic SENP3 by 50%, about the same reduction as achieved by infection with SENP3 shRNA. Moreover, ischaemia did not further decrease in SENP3 levels in the SENP3 knockdown cells. Surprisingly, however, 2h and 18h periods of ischaemia did not significantly reduce SENP3 in the control cells, nor was there and additional reduction in SENP3 levels in the SENP3 knockdown cells. These results indicate that cytosolic levels of SENP3 are rapidly reduced during ischaemia, but that levels recover during prolonged ischaemia.

To examine the link between SENP3 levels and initiation of apoptosis, we tested the effect of SENP3 knockdown on ischaemia-induced cleavage of caspase 3. There were no detectable changes in caspase 3 cleavage following 2h of ischaemia in either control or SENP3 knockdown cells. However, in agreement with the data shown in Fig. 3A, there was a significant increase in cleaved caspase 3 after 2h ischaemia, which was not affected by SENP3 knockdown (Fig. 4). In contrast, cleaved caspase 3 was significantly reduced after 18h ischaemia in both control and SENP3 knockdown cells.

### SENP3 knockdown increases the rate of cell death during ischaemia-reperfusion in H9C2 cells

We next examined how SENP3 affects cell death in response to ischaemia-reperfusion by measuring LDH release. SENP3 knockdown did not alter the total amount of cell death either following 18h ischaemia or 18h ischaemia plus 24h reperfusion (Fig. 5A, B). Interestingly, however, at shorter periods of ischaemia, SENP3 knockdown cells were more susceptible to cell death during ischaemia or ischaemia-reperfusion (Fig. 5 A, B). Indeed, there were significantly higher levels of cell death in the SENP3 knockdown cells sampled after 18h ischaemia and 6h reperfusion compared to cells in which SENP3 levels were not knocked down (Fig. 5C). These data suggest that cells with reduced SENP3 die more quickly during reperfusion following ischaemic insult, supporting a role for SENP3 in cardiomyocyte survival after ischaemia-reperfusion.

**Fig. 5.**
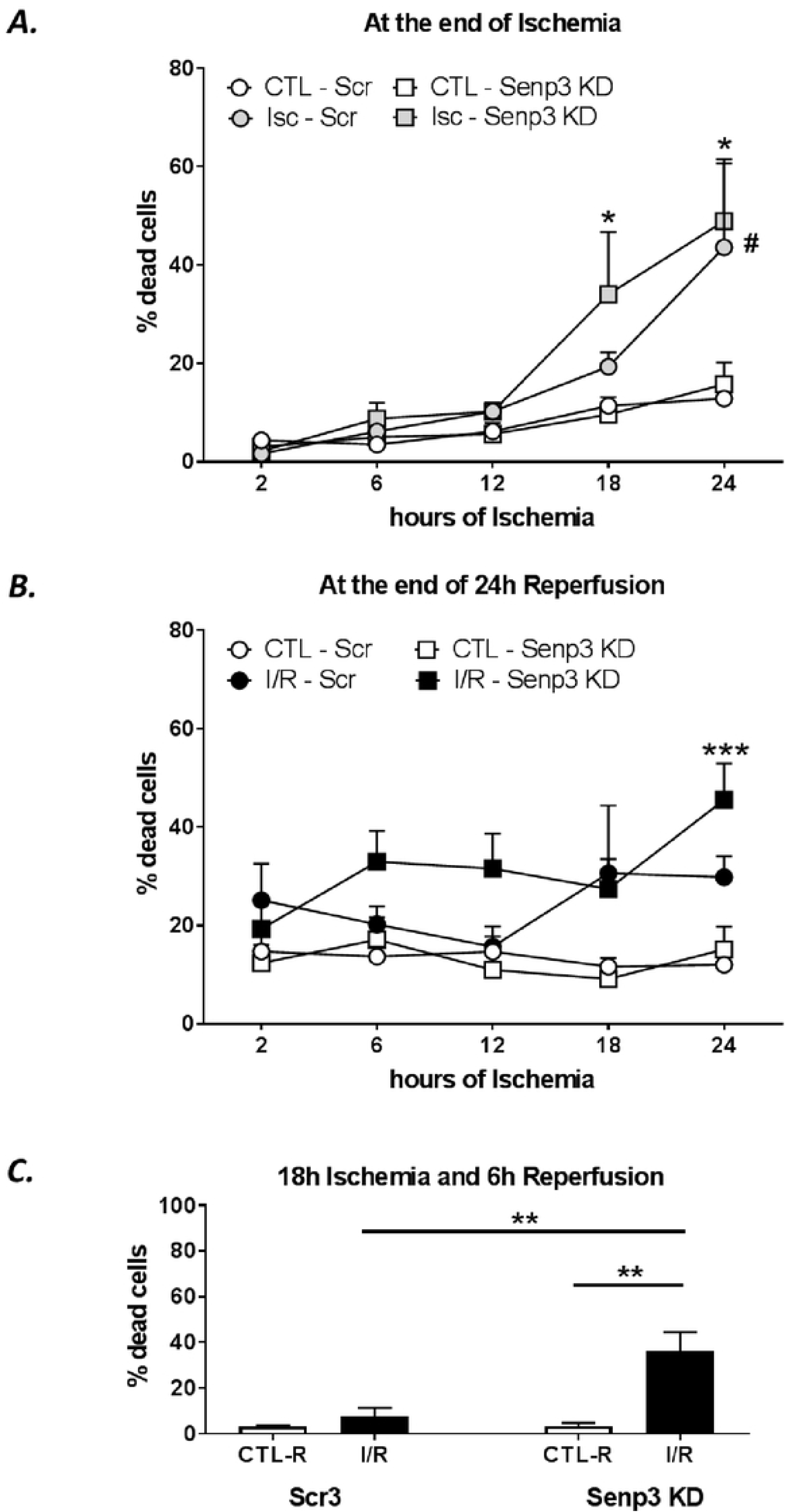
Profile of LDH release after increasing periods of ischaemia in control and SENP3 knockdown H9C2 cells. (**A**) LDH release after indicated times of control (white symbols) and ischaemia (grey symbols), N=3-4 independent experiments. Data presented as mean ± SEM. Quantitative analysis using ordinary two-way ANOVA with Sidak’s correction for multiple comparisons with a pooled variance. * p<0.05 for SENP3 KD group, # p<0.05 for scrambled group, control vs ischaemia. No differences were detected between control and KD cells. (**B**) LDH release after indicated times of control (white symbols) and ischaemia followed by 24h of reperfusion (black), N=3-4 independent experiments. Data presented as mean ± SEM. Quantitative analysis using ordinary two-way ANOVA with Sidak’s correction for multiple comparisons with a pooled variance. *** p<0.001 for SENP3 KD group, control vs ischaemia-reperfusion. No difference between scrambled or KD cells was been observed. (**C**) LDH release after 18h ischaemia followed by 6h reperfusion, N=3 experiments. Data presented as mean ± SEM. Quantitative analysis using ordinary two-way ANOVA test with Sidak’s correction for multiple comparisons with a pooled variance, ** p<0.01.

## Discussion

Increased understanding of the molecular mechanisms underpinning the pathology of cardiac ischaemia and reperfusion is important for the identification of novel drug targets. In particular, strategies to promote the protective changes that occur in preconditioning could lead to cardioprotective therapies for individuals at risk of cardiac ischaemia.

Using total homogenate and subcellular fractions from whole hearts subjected to Langendorff perfusion, we detected some changes in levels of nuclear SUMOylation, and in the SUMOylation of individual mitochondrial substrate proteins. A more striking observation, however, is that levels of the deSUMOylating enzyme SENP3 were dramatically reduced in the cytosol during preconditioning, ischaemia and ischaemia-reperfusion. Moreover, during ischaemia, the decrease in cytosolic SENP3 is paralleled by an increase in nuclear SENP3, suggesting that ischaemia could cause the relocation of SENP3 from the cytosol to the nucleus.

We have previously demonstrated that loss of SENP3 during ischaemia protects HEK293 cells and neurons against reperfusion injury, through promoting SUMOylation of one of its target proteins, Drp1 (21, 25). In heart tissue ischaemia, however, our data suggest a relocation, rather than loss of SENP3. These differences could reflect cell-type specific mechanisms in controlling the localisation, activity and availability of SENP3 during ischaemia. Nonetheless, our data reinforce the concept that SENP3 represents an important stress response protein whose levels and/or localisation are subject to tight control to orchestrate cell survival.

In H9C2 cells ischaemia also leads to a loss of SENP3 from the cytosol. Because these cells are readily amenable to genetic manipulation, we investigated the effects of SENP3 knockdown and show increased susceptibility to cell death following I/R in cells with reduced levels of SENP3. These data are consistent with SENP3 promoting cardiac cell survival after ischaemic stress.

Very recently, two other papers have examined the role of SENP3 in cardiac cell survival after I/R (14, 15). Both of those studies reported an increase in SENP3 levels following I/R in either whole heart or H9C2 cells. We did not observe any changes in SENP3 levels in I/R samples and the reasons for this discrepancy are unclear. However, it remains possible that this is due to differences in the duration or severity of I/R achieved under the different experimental conditions used.

Nonetheless, similar to the observations reported here, Zhang *et al*. observed an increase in I/R-induced H9C2 cell death when SENP3 was reduced. In contrast, Gao *et al*. saw decreased infarct size and improved cardiac function after I/R following SENP3 knockdown in whole heart (14).

Clearly, further work is needed to reconcile these findings, but it remains possible that the role of SENP3 after I/R is dependent on the cell type and population of cells investigated. As a result, the consequences of SENP3 loss in the variety of cell types in whole heart may differ from the effect observed specifically in cardiomyocytes. Indeed, reducing SENP3 promotes HEK293 and neuronal survival after I/R (21, 25), suggesting that the pro-survival role of SENP3 observed here may result from cell type-specific differences in the response to ischaemia.

Consistent with a possible protective role for SENP3 in whole heart, preconditioning prior to ischaemia reduced the extent of SENP3 loss from the cytosol compared to ischaemia alone. Since preconditioning potently protects cells against I/R injury (26), and loss of SENP3 in H9C2 cells promotes cell death upon I/R, these data suggest that the protective effects of preconditioning could, at least in part, be due to the effect of preconditioning in limiting SENP3 loss from the cytosol.

SENP3 deconjugates SUMO2/3 from a wide variety of substrates, the majority of which have not yet been identified (7, 9). Our previous work demonstrated a role for SENP3 in the response to ischaemia through modulation of SUMOylation of its target Drp1 (21, 25), so it is possible that Drp1 is the major SENP3 target mediating its effects in cardiomyocytes. However, our initial experiments suggest that the changes we observed in SENP3 localisation during PCI, ischaemia and I/R are not mirrored by corresponding changes in Drp1 localisation at mitochondria, which is a direct outcome of Drp1 SUMOylation (21, 25, 27). Thus, it seems likely that other, as yet unidentified extranuclear SENP3 targets may be mediating its protective effect on cardiomyocytes. Indeed, it is of note that we observed alterations in specific mitochondria-associated SUMO substrates during PCI, ischaemia and I/R, suggesting an altered SUMOylation profile of a restricted number of cytosolic or mitochondrial substrates may coordinate cardiomyocyte survival after I/R.

Together, our data demonstrate that in whole heart and the H9C2 cardiomyocyte cell line, ischaemia leads to a loss of SENP3 from the cytosol, and suggest that SENP3 loss leads to a reduction in cardiomyocyte survival upon I/R. Thus, promoting SENP3 levels in susceptible individuals, or maintaining the levels of SENP3 after ischaemic infarct, may represent novel therapeutic strategies aimed at promoting cell survival and heart function after cardiac ischaemia.

## Acknowledgements

We thank the BHF and the BBSRC for financial support. We are grateful to Ron Hay (University of Dundee) for the kind gift of the sheep anti-SUMO-1 and anti-SUMO-2/3 antibodies used in this study. We also thank Jordan Martin for assistance in some experiments.

## Author contributions

NR, LL and KAW performed all of the experiments. JMH supervised the project

